# Quantitative Detection of Cell Activity by Measuring the Fluctuation of Intracellular Motility

**DOI:** 10.1101/850602

**Authors:** Morito Sakuma, Yuichi Kondo, Hideo Higuchi

## Abstract

The measurement of cell activity changes during damage is important to understand the process of cell death and evaluate the effect of medicines. To evaluate cell activity generally, we extended the method of intensity fluctuation in which intensity change in the pixel induced by the movement of organelles was calculated. Cancer, endothelial and iPS cells were damaged by reactive oxygen species (ROS) generated by a fluorescent dye (IR700), hydrogen peroxide, and ultraviolet light. The intensity fluctuation in damaged cells gradually decreased independent of the kind of cell, indicating that the decrease in the fluctuation is a general phenomenon in damaged cells. The rupture of vesicles and mitochondria in the cells were observed upon ROS production. The motility of purified kinesin and dynein which transport vesicles and organelles was inhibited by ROS. These suggest that ROS and cytotoxic molecules spreading from ruptured organelles contribute to the reduction in cell activity which brings about the decrease in the motility and intensity fluctuation of organelles driven by kinesin and dynein.

## Introduction

The intracellular condition of cells is changed by external stresses, such as oxidation, no physiological pH and toxins. The accumulation of those stresses leads to cell death (Fujii et al., 2003; Lagadic-Gossmann et al., 2004; Martindale and Holbrook, 2002; Redza-Dutordoir and Averill-Bates, 2016). In the process of cell death, the accumulated damage exceeds a certain threshold, and then the cells switch on the systems for cell death, called apoptosis, or develop coagulative and liquefactive necrosis (Gascoigne and Taylor, 2009; Letai, 2015). These processes are common in various types of cells, including pluripotent stem cells and cancer cells. Therefore, detecting the accumulation of damage is important to evaluate the effect or tolerance of medicines on cells and their culture environment.

Fluorescent probes have been used to detect changes in intracellular conditions during cell death by specifically staining proteins related to cell death (Bussolati et al., 2011; Shi et al., 2012; Yamaguchi et al., 2011; Zhang et al., 1997; Zhang et al., 2015). However, fluorescence probes have problems such as photobleaching and phototoxicity. The accuracy of the detection of cell damage decreases with the decrease in intensity of fluorescent probes induced by photobleaching (Gerlich and Ellenberg, 2003; Laissue et al., 2017; Waters, 2009). Therefore, fluorescent probes for detecting sequential changes in the activity longer than ~10 minutes are not available. Conventional microscopes such as bright field, phase contrast and differential interference contrast microscopes have been used to detect cell activity and damage for a long time because they cause essentially no photobleaching or photodamage (Aftab et al., 2014; Ma et al., 2019; Maddah et al., 2015). In these works, cell activity was usually detected by the morphological and motility changes of whole cells. However, since the morphological changes of cells caused by mild cell damage were not detected and the motility of the cell was stochastic and random, these methods were not precise and quantitative (Aftab et al., 2014; Balvan et al., 2015; Tokumitsu et al., 2010). Therefore, to detect precise and quantitative changes caused by damage over a long time, specific and precise changes in damaged cells should be detected under a conventional microscope.

In previous work, the reduction in the motility of intracellular organelles was detected without morphological changes of cells under a phase contrast microscope (Sakuma et al., 2016). This is supported by the result that the motility of mitochondria was suppressed by cell damage (Debattisti et al., 2017; Liao et al., 2017). The motility of mitochondria gradually decreased just after the addition of hydrogen peroxide, and the decrease in the motility reached 50% within 10 minutes (Debattisti et al., 2017). These results suggested that the effect on the motility of organelles in damaged cells occurred earlier than the changes in cell morphology and motility. Thus, the method of detecting intracellular motility under a conventional microscope would overcome the disadvantages of previous fluorescence, morphological, and cell motility methods. However, little is understood about the generality and mechanism of motility reduction.

In this study, various cells were damaged by several methods, including photoactivation of IR700, irradiation with ultraviolet (UV) light and treatment with hydrogen peroxide (H_2_O_2_), and the change in the motility of organelles was quantitatively evaluated by the developed intensity fluctuation method (IFM). We found that all cells showed a decrease in motility caused by damage. To elucidate the mechanisms of the decrease in motility in damaged cells, we measured the reduction in the motile speeds of single vesicles in cells and in purified kinesin and dynein *in vitro* assays. These measurements revealed that the decrease detected by IMF resulted in the reduction in the organelle motility driven by motor proteins. Since the transport system commonly exists in animal cells, the decrease in the motility or transport of organelles would be a universal phenomenon in various kinds of cells.

## Results

### 1 Decrement of motility of organelles in damaged cells

Changes in the motility of organelles in damaged cells were evaluated by the intensity fluctuation method (IFM) in various cell types. Six kinds of cells, four cancer cell lines, one endothelial cell line, and one iPS cell line, were damaged by the photoactivation of IR700 dye, which is applied for the targeted therapy of cancer cells (Mitsunaga et al., 2011). IR700-EGFP was endocytosed, and the fluorescence of IR700 excited by a red laser was observed inside cells, especially near the nucleus (first and second columns in Fig. 1a). We applied the IFM to detect changes in organelle motility. The changes in intensity fluctuation were shown by heat maps of fluctuation values (third and fourth columns in Fig. 1a). All cells showed a decrease in the fluctuation values induced by photoactivation for one and ten seconds. The mean intensity fluctuation (see Materials and methods) for each kind of cell was calculated (Fig. 1b). The fluctuation values of cells that were illuminated by a red laser without adding IR700 showed no significant changes (left column in Fig. 1b). The fluctuation values in the cancer cells and HUVECs were decreased by ~30% and ~50% at 1 and 10 seconds of photoactivation, respectively, and the corresponding values in iPS cells were decreased by ~15% and ~22% (second and third columns in Fig. 1b). The fluorescence intensity of IR700 in iPS cells was ~10 times lower than that in other cells, indicating that the concentration of IR700 endocytosed in iPS cells was lower than that endocytosed in cancer cells, consistent with the lower decrease in the fluctuation value. In contrast to the significant fluctuation change, the shapes of the cells at 1 minute after 10 seconds photoactivation were not changed (Supplemental Fig. 1c). These results indicated that damage would significantly reduce the motility of organelles without causing significant changes in cell shape.

**Figure 1.**
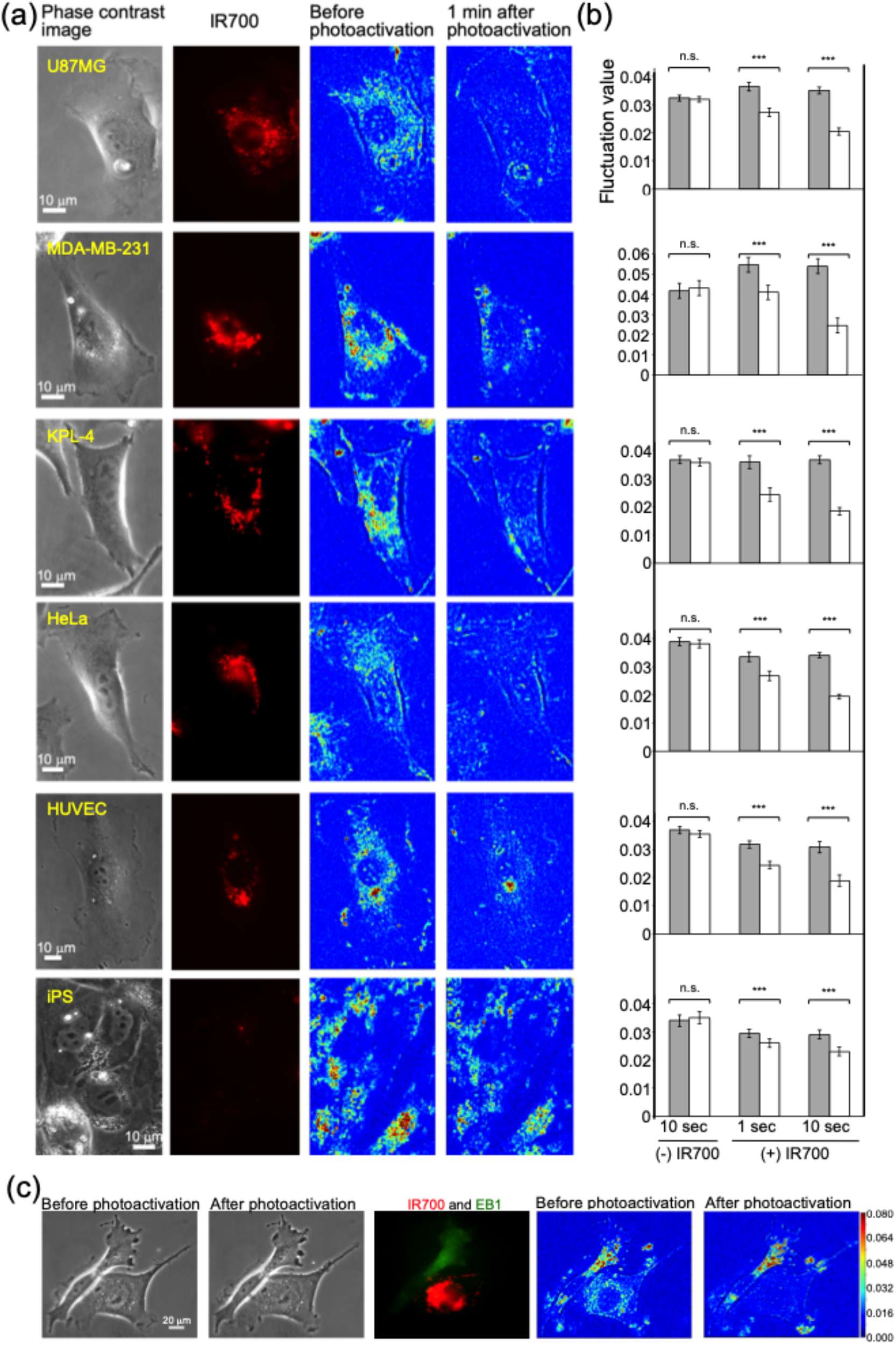
Decreased motility of organelles induced by photoactivation of IR700 in various cell lines. (a) Typical images of the change in motility caused by 10 seconds of photoactivation of IR700 in U87MG, MDA-MB-231, KPL-4, HeLa, HUVEC, and iPS cells. The first column shows phase contrast images before cell damage, and the second column shows the fluorescence of IR700 corresponding to the phase contrast images. The third and fourth columns show heat maps of the change in fluctuation values through photoactivation of IR700 corresponding to the phase contrast images. The scale of the heat maps is the same as that in Fig. 1c. (b) Change in the mean of the fluctuation values with or without IR700 (mean ± SEM) (n.s. denotes not significant, ***p<0.05). IR700 was activated for 1 or 10 seconds. (c) Detection of cell damage in co-cultured U87MG and GFP-EB1-expressing MDA-MB-231 cells. The first and second columns show phase contrast images before and after 10 seconds of photoactivation of IR700. The third column shows the fluorescence of IR700 in U87MG cells and GFP-EB1 in MDA-MB-231 cells. The fourth and fifth columns show heat maps of the change in fluctuation value induced by IR700 photoactivation.

U87MG cells containing IR700 and MDA-MB-231-GFP-EB1 cells without IR700 were cocultured and were illuminated with a red laser (Fig. 1c). Fluorescence images showed that the left and right cells in Fig. 1c were MDA-MB-231 and U87MG cells, respectively (third column in Fig. 1c). Phase contrast images were observed before and at 2 minutes after 10 seconds of photoactivation (first and second columns in Fig. 1c). While the change in cell shape caused by photoactivation could hardly be distinguished in the phase contrast images, the decrease in motility in only U87MG cells after photoactivation was clearly observed. These results indicate that the IFM selectively detected the damaged cells.

Cell damage was also measured by ethidium homodimer-1 (EthD-1), which is one of the conventional methods for detecting cell damage. After 1 second of photoactivation, the nuclei of cells were not stained significantly by EthD-1 (Supplemental Fig. 2). The nucleus was stained at 1 hour but not 30 minutes after photoactivation, suggesting that the IFM detected cell damage more rapidly than staining with EthD-1.

**Figure 2.**
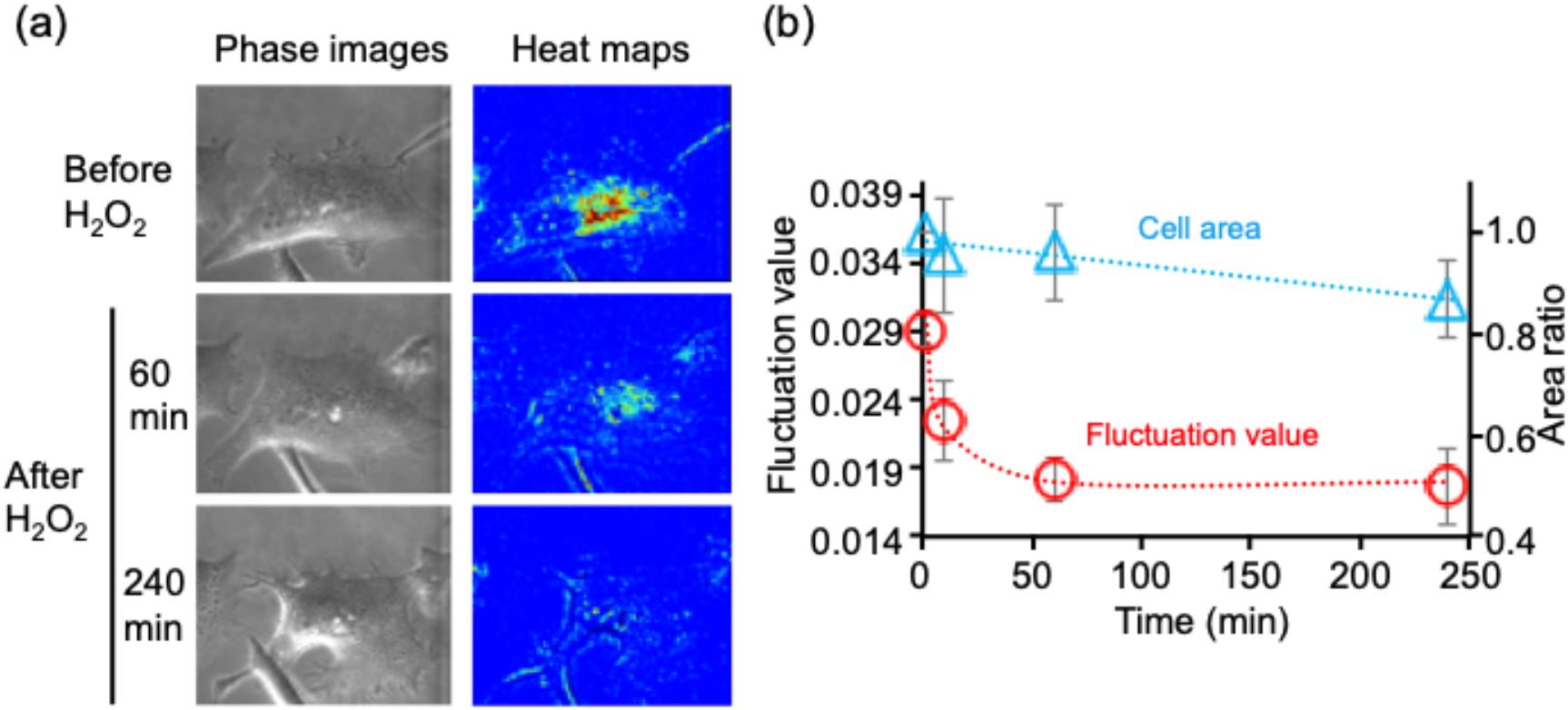
Decreased motility of organelles induced by the addition of hydrogen peroxide (H_2_O_2_). (a) Typical phase contrast images of cell and heat maps of fluctuation values corresponding to the phase contrast images before and after the addition of H_2_O_2_. (b) Change in fluctuation value (red circles) and cell area (light blue triangles) caused by the addition of H_2_O_2_ (mean ± SEM). The first and second y-axes denote the mean fluctuation values and area ratios, respectively. Broken lines show exponentially fitted lines.

### 2 Motility changes of organelles in oxidative stress

To evaluate whether the decrease in motility could become an indicator of cell damage, cells were damaged by hydrogen peroxide (H_2_O_2_) treatment and ultraviolet (UV) light irradiation. H_2_O_2_ was added to the culture medium at a final concentration of 100 μM, and phase contrast images of cells were observed for 4 hours (Fig. 2a). In the phase contrast images, apparent changes in size and shape could hardly be observed within 60 minutes. At 240 minutes, needle-like structures were observed near the edge of cells that maintained their desmosomes. The cell area gradually decreased until 240 minutes (slope: −0.05 (% area/minutes); blue triangles in Fig. 2b). The fluctuation value decreased exponentially with the time over 10 minutes (red circles in Fig. 2)). A decrease in the fluctuation was also observed after UV irradiation without a significant change in cell area (Supplemental Fig. 3). These results indicated that the intensity fluctuation was decreased by oxidative stress and that the decrease in the intensity fluctuation was more rapid than the changes in the cell area.

### 3 Ruptures of vesicle and mitochondria by photoactivation of IR700

To understand the reason for the reduction in the motility of organelles, the changes in the intercellular organelles in damaged cells were measured. At 1 hour after adding IR700, most of the IR700 fluorophores were observed near the nucleus (Fig. 3a). The fluorescence intensity of those vesicles gradually decreased upon irradiation with the red laser because of photobleaching and then decreased rapidly at 3-4 seconds (Fig. 3a and b). Diffusion of IR700 fluorescence from the large vesicles to the cytoplasm was also observed (Supplemental Fig. 4), indicating that vesicles containing IR700 became ruptured or leaky and that IR700 and toxic molecules, such as protease and reactive oxygen species, would diffuse in the cytosol.

**Figure 3.**
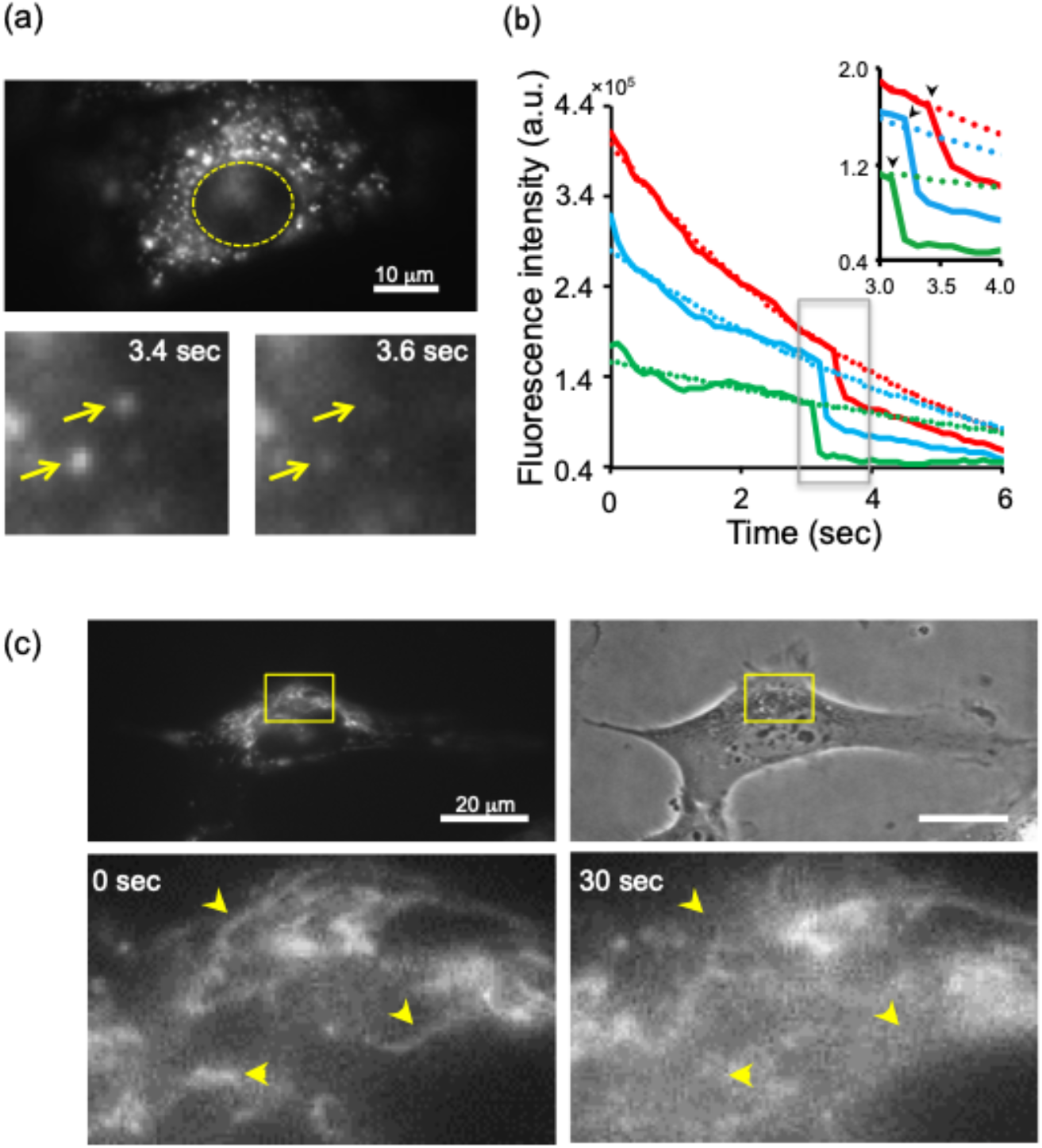
Real-time imaging of damage to organelles induced by the photoactivation of IR700. (a) Real-time imaging of IR700 fluorescence in vesicles. The upper image shows the distribution of the fluorescence of IR700. The yellow broken line shows the periphery of the nucleus. The lower images show the rapid decrease in IR700 fluorescence in vesicles (yellow arrows). (b) Trajectories of the fluorescence intensity of IR700. Solid and broken lines show the change in measured fluorescence intensity and exponentially fitted lines within 2.5 seconds, respectively. Black arrows in the inset image show the rapid decrease in the intensity caused by IR700 photoactivation. (c) Dynamics of mitochondria upon photoactivation of IR700. Upper images show mitochondria stained by the fluorescent probe CellROX (left) and the corresponding phase contrast image (right). Lower images show changes in the distribution of mitochondria upon photoactivation for 30 seconds. Lower images are the magnified views of the areas in the white rectangles in the upper images. Yellow arrows indicate that the rod-shaped structure of mitochondria disappeared upon photoactivation of IR700.

Mitochondria are one of the organelles that could be observed by a phase contrast microscope. Next, mitochondria were stained with CellROX, which is an indicator of reactive oxygen species (ROS), to observe their movement following cell damage. Mitochondria generate ROS and thus can be stained by CellROX (left and upper panels in Fig. 3c). Under photoactivation, the tubule-like structure of the mitochondria rapidly disappeared, and the fluorescence of IR700 did not accumulate in the mitochondria (right and lower panel in Fig. 3c). At the same time, the fluorescence intensity in the cytoplasm was increased, indicating that ROS in the mitochondria diffused into the cytoplasm by mitochondrial rupture, thereby enhancing the fluorescence intensity of CellROX in the cytoplasm (right and lower panels in Fig. 3c). These results suggested that vesicles and mitochondria were damaged by the photoactivation of IR700, and toxic molecules, including proteases and ROS, diffused from ruptured organelles to the cytoplasm.

### 4 Motility of vesicles in damaged cells detected by quantum dots

In phase contrast microscopy, various shapes and sizes of organelles were observed. The IFM mainly detected changes in the motility of organelles near the nucleus, where endosomes and lysosomes are mainly located. Thus, the change in those organelles was directly observed and evaluated by single-particle tracking of QD-EGFR conjugates. QD-EGFR was endocytosed in vesicles and transported near the nucleus (Fig. 4a). Before damaging the cell, the vesicle-QDs moved linearly and randomly (blue line in the inset image of Fig. 4b). The linear region is presumably transported by molecular motors such as kinesin and dynein. After cell damage, the motility of the vesicles was decreased, and the trajectories of the vesicles-QDs showed confined movement (red line in the inset image). The mean-square displacements (MSDs) were also decreased by cell damage. The diffusion coefficients before and after damage were obtained by fitting the MSD plot with the equation *4Dt*, where *D* and *t* are the diffusion coefficient and time, respectively. The diffusion coefficients obtained before and after the damage were 0.42 μm^2^/s and 0.07 μm^2^/s, respectively. These results indicated that the diffusion after damage was approximately 6 times lower than that before damage.

**Figure 4.**
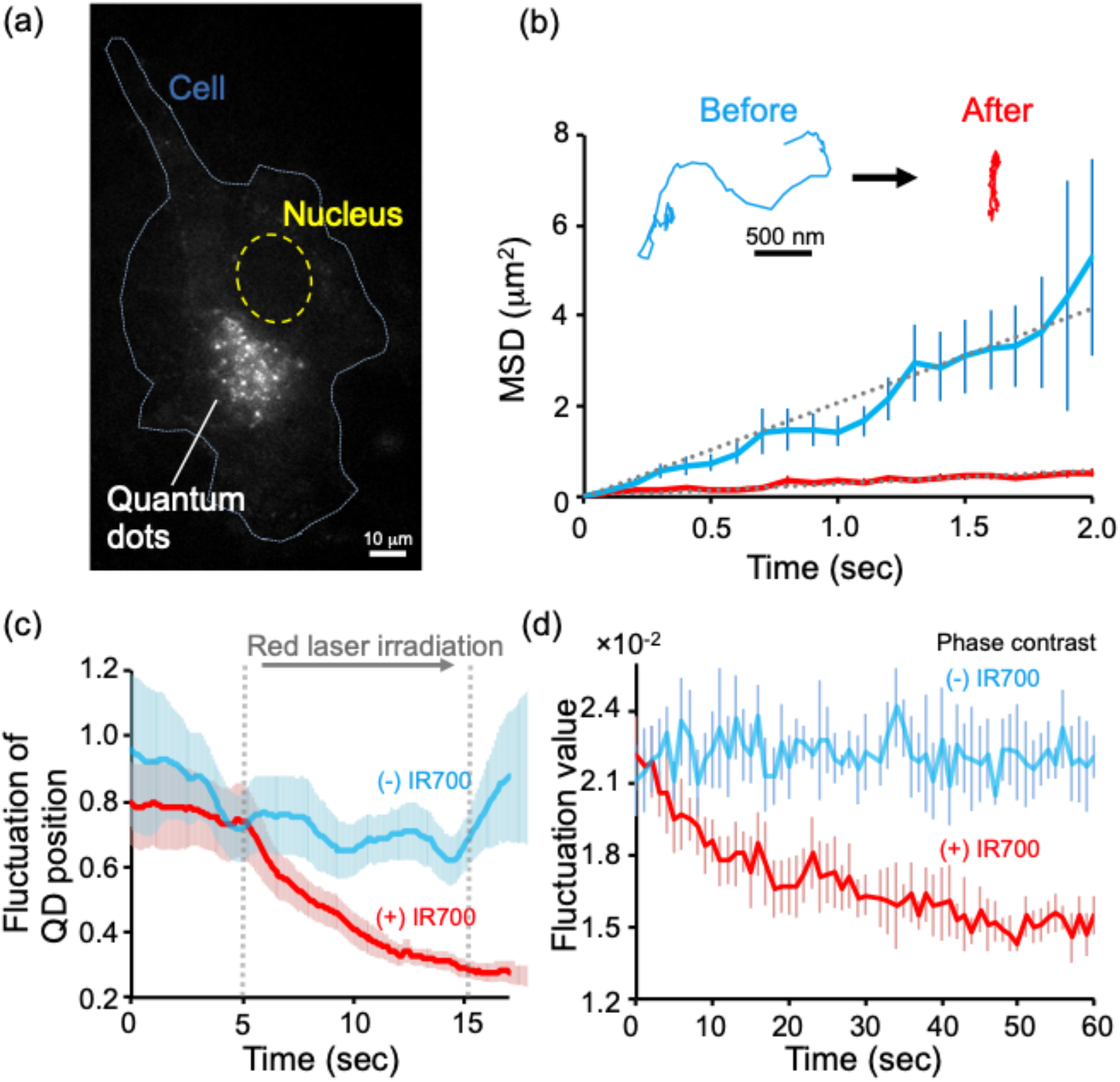
Single-particle tracking of vesicles labeled with quantum dots (vesicle-QDs) in damaged cells. (a) Distribution of vesicle-QDs inside cells. Blue and yellow broken lines show the periphery of the cell and nucleus, respectively. (b) Mean-square displacement (MSD) plot of the motility of vesicle-QDs before and after IR700 photoactivation (mean ± SEM). Light blue and red solid lines show the MSD before and after photoactivation, respectively. Inset image shows a typical trajectory of vesicle-QDs. Broken gray lines show lines linearly fit to the MSD values. (c) Fluctuation of the position of vesicle-QDs upon photoactivation with or without using IR700 (mean ± SEM). The cells were irradiated with a red laser from 5 to 15 seconds (gray broken line). (d) Real-time change in fluctuation values calculated from phase contrast images with or without using IR700 (mean ± SEM). The cells were irradiated with a red laser from 2 to 32 seconds.

The timing of the decrease in vesicle QDs was evaluated in damaged cells and compared with the results of the IFM. The fluctuation of the position of the vesicles-QDs was measured instead of the intensity fluctuation since the fluorescence intensity of QDs gradually changed upon excitation with a green laser and photoactivation of IR700 and since the fluctuation of QDs could hardly be precisely evaluated (Supplemental Fig. 5) (Li et al., 2006). The fluctuation of position was calculated from the mean standard deviations of the displacement of vesicles-QDs at every 10 frames (Fig. 4c). The fluctuation of position gradually decreased after photoactivation of IR700. The time-course change of the intensity fluctuation under a phase contrast microscope was also measured, and the decrease was compared between vesicles-QDs and IFM (Fig. 4d). The intensity fluctuation of organelles gradually decreased in parallel with the decrease in the fluctuation of the position of the vesicles-QDs, suggesting that this change in the motility of endosomes and lysosomes was coupled with the change in the motility of organelles detected by the IFM.

### 5 Decrease in motilities of kinesin and dynein by oxidative stress

Vesicles including endosomes and lysosomes are transported by molecular motors, such as kinesin and dynein, along with microtubules. Therefore, the effects of IR700 photoactivation and hydrogen peroxide (H_2_O_2_) treatment on motor proteins were evaluated by an *in vitro* motility assay (Fig. 5a) (Higuchi et al., 2002). Bovine serum albumin (BSA) conjugated with IR700 and biotin were bound to the surface of a glass chamber. Biotinized kinesin or dynein was attached to BSA-biotin via streptavidin, and then microtubules were flowed into the chamber (Fig. 5a). The microtubules were distributed homogeneously before photoactivation (left column in Fig. 5c). After photoactivation, the microtubules accumulated within the area of photoactivation as a result of slow movement, while the motility outside the area was hardly changed (right column in Fig. 5c). The position of the leading edge of the microtubules was tracked every 10 seconds to understand the velocity change in response to oxidative stress (Fig. 5b and d). Microtubules driven by kinesin and dynein moved linearly and continuously before damage, while the motility decreased very much inside the red circles (Fig. 5d). The velocity after the photoactivation of IR700 and addition of H_2_O_2_ was analyzed from the trajectories of microtubules inside the photoactivated area (Fig. 5b). After photoactivation, the mean velocity of the microtubules driven by kinesin and dynein was decreased from 394 to 6 nm/s and from 99 to 4 nm/s, respectively. The velocity also decreased from 467 to 48 nm/s at 60 minutes in the presence of 0.1 mM H_2_O_2_. It is noted that the motility of kinesin was not inhibited by red laser irradiation without using IR700 (Supplemental Fig. 6a). The addition of an ROS scavenging system (GCOβ) also prevented the decrease in the motility of kinesin caused by the photoactivation of IR700 (Supplemental Fig. 6b). These results indicate that the decrease in the motility of kinesin and dynein was induced by oxidative stress.

**Figure 5.**
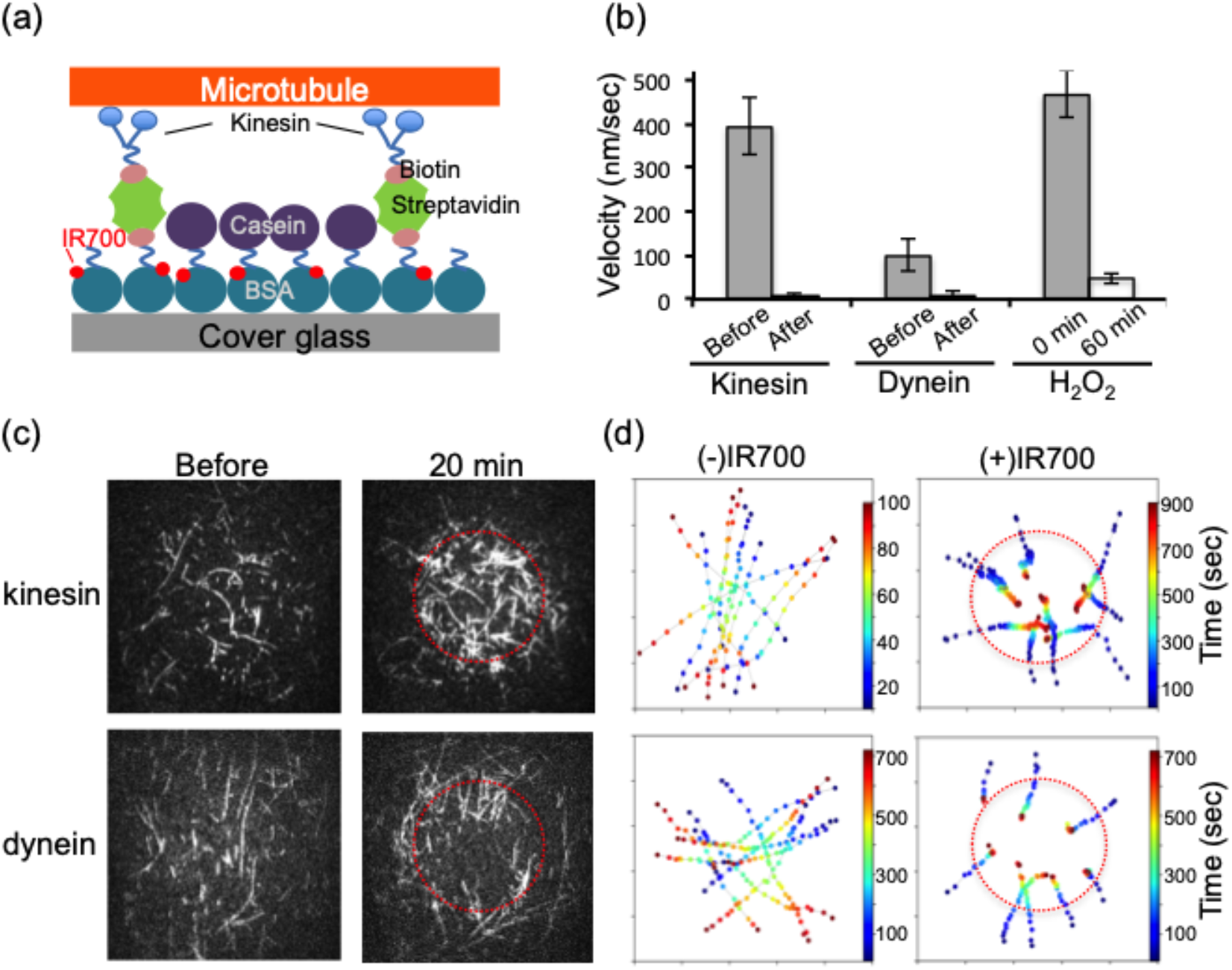
Motility assay of kinesin and dynein on microtubules under oxidative stress. (a) Schematic illustration of the motility assay. Microtubules were fixed on the cover glass by binding with kinesin or dynein. IR700 binds to BSA through the NHS moiety. Photoactivation of IR700 damages molecular motors and microtubules. (b) Decreased motility of kinesin and dynein upon photoactivation of IR700 and addition of hydrogen peroxide (H_2_O_2_). Gray and white bars indicate the velocity before and after photoactivation and addition of H_2_O_2_ (mean ± SEM), respectively. The red laser was irradiated for 100 milliseconds, and the final concentration of H_2_O_2_ was 100 μM. (c) Change in the distribution of microtubules caused by photoactivation. The red laser was irradiated during the period inside the red broken lines, and the fluorescence of the microtubules was observed for 20 minutes. (d) Trajectories of microtubules before and after photoactivation. Color bars indicate observation time, and red broken lines denote the spots irradiated with the red laser corresponding to (c).

## Discussion

### Advantage of the intensity fluctuation method (IFM) in detecting cell damage

The motility of organelles in several cell lines, including cancer cells, endothelial cells and iPS cells, was detected by the intensity fluctuation method (IFM) (Fig. 1a). The sensitivity and quantitativity of measurements of the fluorescence probes decrease by the photobleaching, phototoxicity and background of the free probes (Laissues et al., 2017; Waters 2009). Actually, a background of EthH-1 in the cytoplasm and nucleus was high even in intact cells, and the fluorescence intensity in the nucleus showed diversity among each cell (Supplemental Fig. 2). The IMF detected damage of cells even when EthH-1 staining did not detected the damage (Sakuma et al. 2016).

Observation of cell morphology and motility under label-free microscopes such as phase contrast is not so quantitative because of polymorphism of cell (Ebara et al., 2018; Li et al, 2006; Masuzzo et al., 2016). In this study, the damage of cell was detected by IFM even when the changes in the areas and shapes of the cell were not detected (Fig. 2 and Supplemental Fig. 1c). These indicate that the IFM is more sensitive and quantitative than the fluorescence, motility and morphology methods (Fig. 1, 2, and 4).

### Oxidative stresses induced the decrement of the motility of organelles

Intensity fluctuation that indicates the motility of organelles was gradually decreased upon IR700 photoactivation, H_2_O_2_ treatment, and UV irradiation. What material generated from these methods suppressed the motility of organelles? Photoactivation of IR700 was recently developed to specifically damage cancer cells *in vitro* and *in vivo* (Harada et al., 2015; Nagaya et al., 2016). Previous research has shown that photoactivation induces cellular damage by producing ROS and heat (Mitsunaga et al., 2011; Kishimoto et al., 2015). To specify the effect of photoactivation, we applied a motility assay of kinesin and dynein to microtubules (Fig. 5). The motility of microtubules inside the photoactivated region rapidly decreased (Fig. 5b-d). However, the motility of kinesin in the presence of oxygen scavenging reagents that removed ROS was almost constant upon IR700 photoactivation (Aitken et al., 2008; Harada et al., 1990) (Supporting Fig. 6b). Therefore, our results indicated that the production of ROS induced by the photoactivation of IR700 is a main factor in inhibiting the motility of motor proteins. H_2_O_2_ is one form of ROS and indiscriminately induces oxidation stress toward protein, lipids, mitochondria, and DNA (Redza-Dutordoir and Averill-Bates, 2016). UV irradiation also produces ROS from cytoplasmic molecules and damages proteins and DNA (Heck et al., 2003). Therefore, all methods produced ROS for damaging cells, and the decrease in the motility of organelles driven by motor proteins would be an indicator of cell damage induced by ROS.

### Universality of the decrease in the motility of organelles in damaged cells

The intensity fluctuation detected the movement of organelles including vesicles, mitochondria, Golgi and endoplasmic reticulum. Most animal cells showed the dynamics of those organelles, and thus, our IFM could easily be applied to detect damage in various types of cells.

We showed the inhibition of organelle transportation at high levels of ROS. This result is consistent with the finding that an elevated level of ROS induced by mitochondrial and lysosomal damage is observed during apoptosis in many cell types (Murphy MP, 2013; Redza-Dutordoir and Averill-Bates, 2016). Not only ROS but also acid and protease leaked from the damaged mitochondria and lysosomes would inhibit organelle transportation (Boya and Kroemer, 2008). Acidification of the cytoplasm is also observed in the process of apoptosis and with increases in ROS (Clément et al., 1998; Lagadic-Gossmann et al., 2004, Sakuma et al., 2016). The bacterial cytoplasm showed decreased diffusion at lower cytoplasmic pH (Maharana et al., 2016), and thus, the change in physical properties of the cytoplasm would correlate with the decrease in the motility of organelles. Therefore, a decrease in organelle motility is generally observed in various types of cell damage and cell death.

## Conclusion

Our novel intensity fluctuation method (IFM) analyzing phase contrast images quantitatively detected cell damage. The reduction in the motility of organelles in damaged cells was successfully measured by the decrease in intensity fluctuation. Vesicle transport in damaged cells and the motility of purified motor proteins were inhibited by cell damage, indicating that a reduction in the motility of motor proteins and organelle transport would induce a decrease in the intensity fluctuation. The reduction in the motility of organelles was also observed upon oxidative stress induced by ultraviolet light irradiation and H_2_O_2_ treatment. Reactive oxygen species would be produced in the process of cell death, and thus, the decrease in motility would be commonly observed in damaged cells. Therefore, our IFM would be applicable to the facile and quantitative detection of several kind of cell damage or cell death.

## Materials and method

### Cell culture and preparation

Glioblastoma cells (U87MG, American Type Culture Collection, Manassas, Virginia), breast cancer cells (KPL-4, Kawasaki Medical School) (Tada et al., 2007), and cervical cancer cells (HeLa, Cell Resource Center for Biomedical Research·Cell Bank at Tohoku University) were cultured at 37 °C with 5% CO_2_ and 95% air in Dulbecco’s modified Eagle’s medium (DMEM, Sigma-Aldrich, St. Louis, Missouri) supplemented with 10% fetal bovine serum (FBS, Takara Bio Inc., Shiga, Japan), 1% L-glutamine (Nacalai Tesque, Kyoto, Japan), and 100 U/mL penicillin-streptomycin (Wako Pure Chemicals Industries, Osaka, Japan). Wild-type MDA-MB-231 (Summit Pharmaceuticals International) and GFP-EB1 (end binding protein-1)-expressing MDA-MB-231 cells were cultured at 37 °C without CO_2_ in Leibowitz’s L-15 medium (L-15, ThermoFisher Science, Waltham, Massachusetts) supplemented with 10% FBS, 1% L-glutamine, and 100 U/mL penicillin-streptomycin. Human umbilical vein endothelial cells (HUVECs, Thermo Fisher Science) were cultured at 37 °C with 5% CO_2_ and 95% air in Medium 200 (Thermo Fisher Science) supplemented with 50× low serum growth supplement (LSGS, Thermo Fisher Science) and 100 U/mL penicillin-streptomycin. Human induced pluripotent stem cells (iPS, The Institute of Medical Science, The University of Tokyo) were cultured at 37 °C with 5% CO_2_ and 95% air in ReproFF (ReproCELL, Kanagawa, Japan) supplemented with 5 ng/mL basic fibroblast growth factor (bFGF, PeproTech Inc., Rocky Hill, New Jersey). U87MG, KLP-4, HeLa, MDA-MB-231, and HUVECs were confluently cultured on polystyrene cell culture dishes and recovered by using 1× TrypLE express enzyme (Thermo Fisher Science). The concentration of recovered cells was adjusted to 1.0 × 10^4^ cells/mL, and the suspension was seeded on a glass-bottom dish (Matsunami Glass Ind., Ltd., Gunma, Japan) coated with 2% collagen. iPS cells were cultured on cell culture dishes coated with a Matrigel (Corning, Tewksbury, Massachusetts) and formed colonies. After the size of the colonies reached approximately 1 mm, the cells were treated with dissociation solution for human ES/iPS cells (ReproCELL) and recovered by using a cell scraper. The colonies were broken up to approximately 200 μm by gentle pipetting and seeded on a glass-bottom dish coated with Matrigel. After being cultured for one day at 37 °C, all cells were prepared for the following experiments.

### Fluorescence staining of cells

Mitochondria in cells were stained by CellROXorange (Thermo Fisher Science). The final concentration of CellROX was 5 μM, and the fluorescence of CellROX was observed after incubating for 10 minutes at 37 °C and washing. Quantum dots (QD605) were used to observe the movement of vesicles in cells. QD605 was conjugated with an antibody against epithelial growth factor receptor antibody (Abcam, Cambridge, United Kingdom) (QD-EGFR) by using an antibody conjugation kit (Qdot^™^ 605 Antibody Conjugation Kit, Thermo Fisher Science). QD-EGFR was added to the culture medium of cells at a final concentration of 5 nM, and the cells were incubated for 10 minutes at 37 °C. After washing three times with culture medium and incubating for 1 hour at 37°C, the fluorescence of endocytosed QD-EGFR was observed. Cell damage was evaluated by ethidium homodimer-1 (EthD-1). EthD-1 was added at a final concentration of 5 μM after damaging cells.

### Damaging cells

Cells were damaged by the photoactivation of IR700, addition of hydrogen peroxide (H_2_O_2_), and irradiation with ultraviolet (UV) light. IRDye 700DX NHS ester (IR700, Li-COR Bioscience, Lincoln, Nebraska) and anti-EGFR antibody (Abcam) were conjugated for 2 hours at room temperature in 10 mM Na_2_HPO_4_ (pH 8.0) (Mitsunaga et al., 2011). The conjugates (IR700-EGFR) were added to the culture medium of cells at a final concentration of 1 μM, and the cells were incubated for 1 hour at 37 °C and washed two times with culture medium. To damage cells by oxidative stress, the cells were treated with hydrogen peroxide (H_2_O_2_, Wako Pure Chemicals). H_2_O_2_ was added to the culture medium of cells at a final concentration of 100 μM. After the addition of H_2_O_2_, the cells were continuously observed by a phase contrast microscope. Cells were damaged by ultraviolet (UV) light at a wavelength of 360-370 nm including a Hg-lamp spectrum. A fluorescence filter set (excitation filter: BP360-370, emission filter: BA420, dichromatic mirror: DM400, U-MNUA2, Olympus, Tokyo, Japan) and a mercury lamp (Olympus) were used for UV light irradiation, and the cells were irradiated with UV light for 10 seconds.

### Fluorescence microscopy for observing cells and proteins

All images of cells were taken by an inverted microscope (IX-70, Olympus) equipped with a phase contrast objective lens (PLAPON 60XOPH, Olympus), halogen and mercury lamps (U-LH100 and U-ULS100HG, Olympus), EMCCD cameras (iXon 3, Andor Technology Ltd., Belfast, Northern Ireland), and an incubator (TOKAI HIT Co., Ltd., Shizuoka, Japan) as reported previously with modifications (Sakuma et al., 2016). The temperature and concentration of CO_2_ were maintained at 37 °C and 5%, respectively, by the incubator. During the acquisition of phase contrast images, halogen light was passed through a 510-550 nm bandpass filter to prevent photoactivation of IR700. IR700 was photoactivated by using a red laser (635 nm, 100 mW, Barrington, Edmund Optics, New Jersey), and the fluorescence was observed by a camera equipped with a 690-730 bandpass filter. The fluorescence of QD-EGFR, CellROX, rhodamine, and EthD-1 was observed with illumination by a green laser (532 nm, Showa Optronics, Tokyo, Japan) and detected by a camera (EMCCD, Andor iXon-plus 885) equipped with a 600-620 nm bandpass filter. GFP-EB1 was observed under a blue laser (488 nm, Showa Optronics) and detected by a camera equipped with a 510-550 nm bandpass filter.

### Intensity fluctuation method (IFM) to detect cell damage

To detect the motility change of organelles caused by cell damage, the intensity fluctuation of pixels in phase contrast images was calculated. The intensity fluctuation (ΔI/<I>) was calculated by the standard deviation (ΔI) of the intensity divided by the mean intensity (<I>) of a pixel according to previous methods with modification (Sakuma et al., 2016). The phase contrast images of cells were taken for 300 frames at 10 (Fig. 4d) or 20 (other figures) frames/second. To enhance the movement of organelles, the phase images were processed by a bandpass filter with a bandwidth of 3-10 pixels (390-1300 nm) included in ImageJ software (Sakuma et al., 2016). Then, all phase contrast images were processed with 4 × 4 binning to reduce the number of pixels in order to reduce the intensity noise and save calculation time. The intensity fluctuation was calculated every 10 frames (1 and 0.5 seconds). A total of 6 rectangles in each cell with a size of 36 (6 × 6) binned pixels were selected randomly near the nucleus (Supplemental Fig. 1a). The mean intensity fluctuation of 36 binned pixels was defined as one set of intensity fluctuations. The p-value of fluctuation values between damaged and non-damaged cells from Student’s t-test was 0.07±0.04 (mean ± SE for 4 cells) at one set (10 frames in Supplemental Fig. 1b; p-value was out of range). To reduce the p-value, the mean ± SE of multiple sets of fluctuation was calculated. The p-value decreased with increasing number of sets (or frames) (green rectangles in Supplemental Fig. 1b). To ensure a significant difference between the damaged and non-damaged cells, we took the mean of 30 sets (or 300 frames) as the intensity fluctuation. Custom code in Python was used for the calculation and creation of heat maps.

### Single-particle tracking of vesicles by using quantum dots

Vesicles inside cells were labeled by quantum dot-EGFR conjugates (QD-EGFR). The fluorescence of vesicles was observed for 17 seconds at 10 frames/s including the photoactivation period (10 seconds) of IR700. The localization of vesicles was estimated by fitting fluorescence-intensity profiles of QDs with a two-dimensional Gaussian function (Yildiz and Selvin, 2005). From the trajectories, the ensemble-averaged mean-square displacement (MSD) was calculated by the following equation:

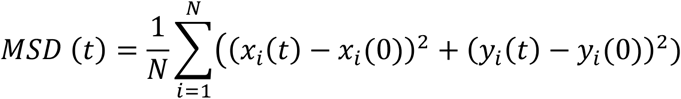

where *N* is the number of particles and *x_i_*(*t*) and *y_i_*(*t*) are the coordinates of particle *i* at time *t*. In total, 32 and 29 particles with and without IR700, respectively, were tracked, and then the MSD was calculated on each trajectory. The mean MSD from t = 0 to 2 seconds was calculated and fitted to a linear equation for estimating the diffusion coefficient (D) using the equation *MSD* = *4Dt* (Maharana et al., 2016).

The fluctuation of the localization of QDs was calculated from the standard deviation of the displacement of the QDs. The standard deviation of the displacement in 10 frames was calculated and then shifted by one frame in a similar manner to a moving average through a total of 170 frames.

### Motility assay of kinesin and dynein

A motility assay of kinesin and dynein was performed on cover glass in a flow chamber (Fig. 5a). First, 1 mg/mL bovine serum albumin (BSA) conjugated with biotin was flowed to cover the surface of the glass. Then, IR700 was flowed to conjugate with the BSA through the NHS moiety. Next, 1 mg/mL streptavidin was flowed to bind with biotin, and then 0.5 mg/mL casein was flowed to prevent the nonspecific binding of kinesin and dynein to BSA or the glass surface. Dimer kinesin (mouse KIF5A with 490 amino acid) or dynein (human dynein-1, GST-D384) labeled with BDTC were flowed to bind with streptavidin, and tubulin flowed. Purification of kinesin, dynein, and microtubules and labeling of the microtubules by rhodamine were carried out according to previous reports (Kinoshita et al., 2018). By adding 1 mM of ATP, the gliding of microtubules labeled with rhodamine was observed by fluorescence microscopy

## Acknowledgements

We thank Grant-in-Aid for Scientific Research on Scientific Research (A) and (B) (H.H. 23247022, 16H04773) from the Japan Society for the Promotion of Science and Scientific Research (C) (S.K. 26440075)

