## Supplemental Figures for "Quantitative Detection of Cell Activity by Measuring the Fluctuation of Intracellular Motility"

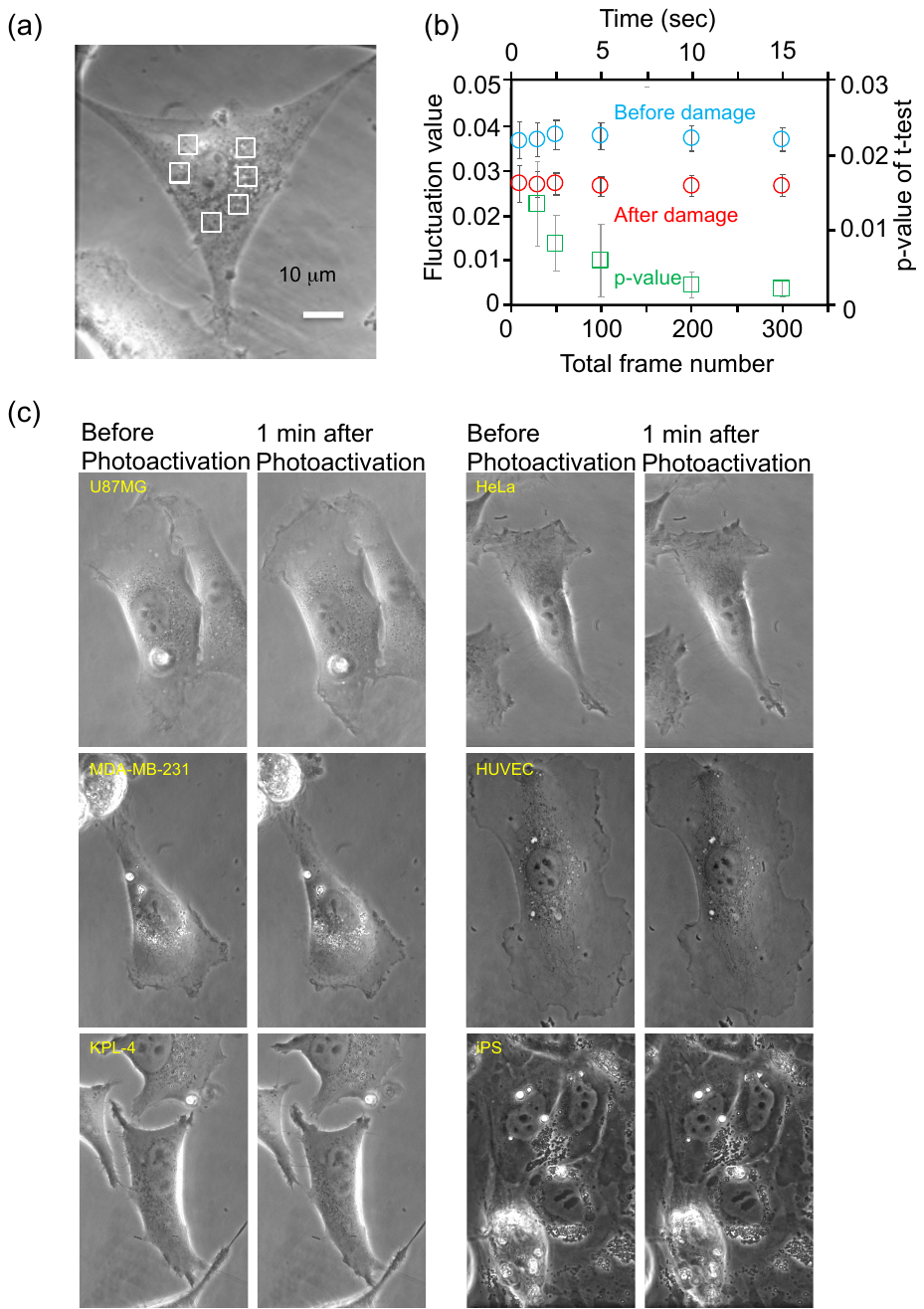

**Supplemental Figure 1.** Setting for measuring the fluctuation value caused by cell damage with the intensity fluctuation method (IFM). (a) ROI sets for detecting fluctuation values. White rectangles (25 × 25 pixels) show the detection area of the intensity fluctuation value. (b) Effect of total frame number on accuracy of damage detection by the IFM. The first and second x-axes indicate the total frame number and corresponding time for measurement, and the first and second y-axes show the fluctuation values before and after cell damage (light blue and red circles) and the p-value of the t-test corresponding to the fluctuation values (mean ± SD), respectively. The p-value at 10 frames was outside the range (0.07±0.04 (mean ± SE)). Fluctuation values were calculated from four U87MG cells. (c) Change in the shape of cells upon photoactivation of IR700. Left and right images show phase contrast images before photoactivation and at 1 minute after photoactivation, respectively.

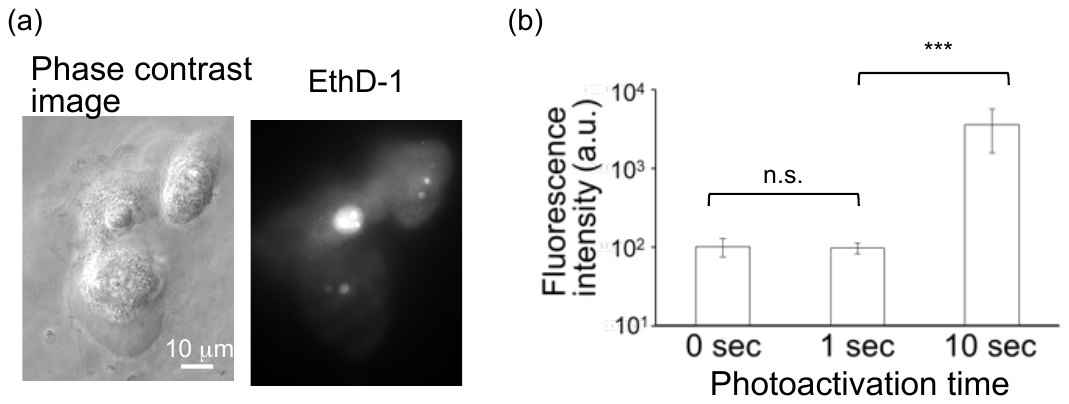

**Supplemental Figure 2.** Detection of cell damage by using ethidium homodimer-1 (EthD-1). (a) Phase contrast image of and the fluorescence of EthD-1 in U87MG cells damaged by the photoactivation of IR700. (b) The fluorescence intensity of the nucleus stained with EthD-1 (mean ± SEM). IR700 was photoactivated for 0, 1, and 10 seconds, and the fluorescence intensity of EthD-1 was measured 1 hour after photoactivation (n.s. denotes not significant, ***p<0.001).

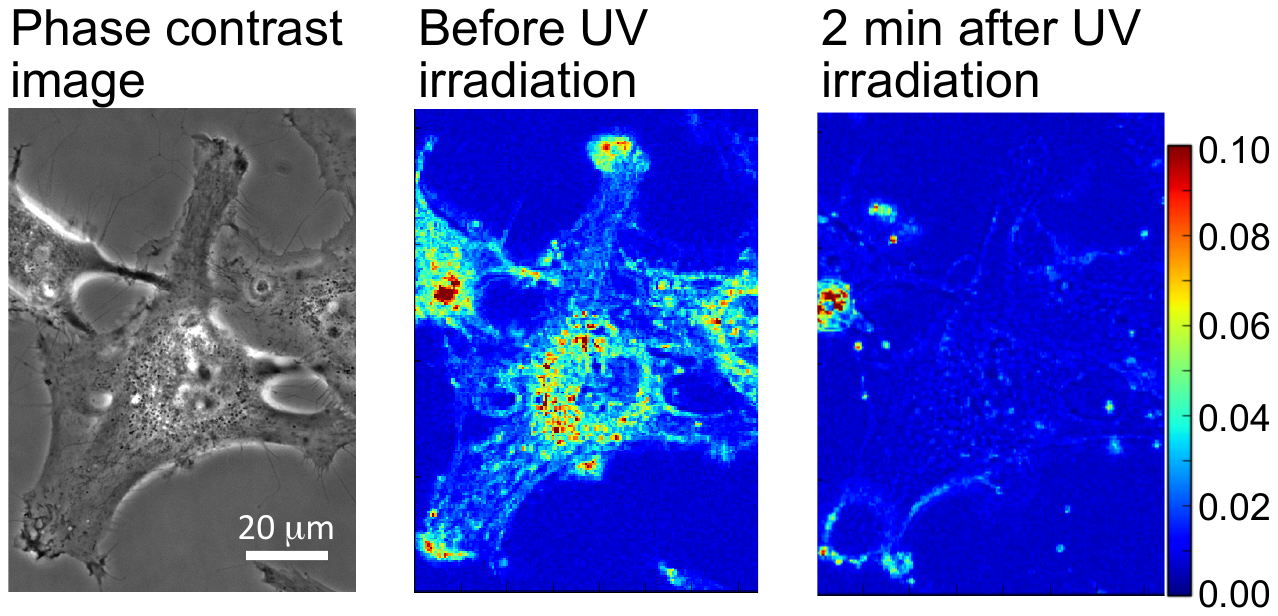

**Supplemental Figure 3.** Change in fluctuation value upon ultraviolet (UV) light irradiation. The left image shows a phase contrast image before UV irradiation, and the center and right images show heat maps of the fluctuation value before and after UV irradiation, respectively, corresponding to the phase contrast image.

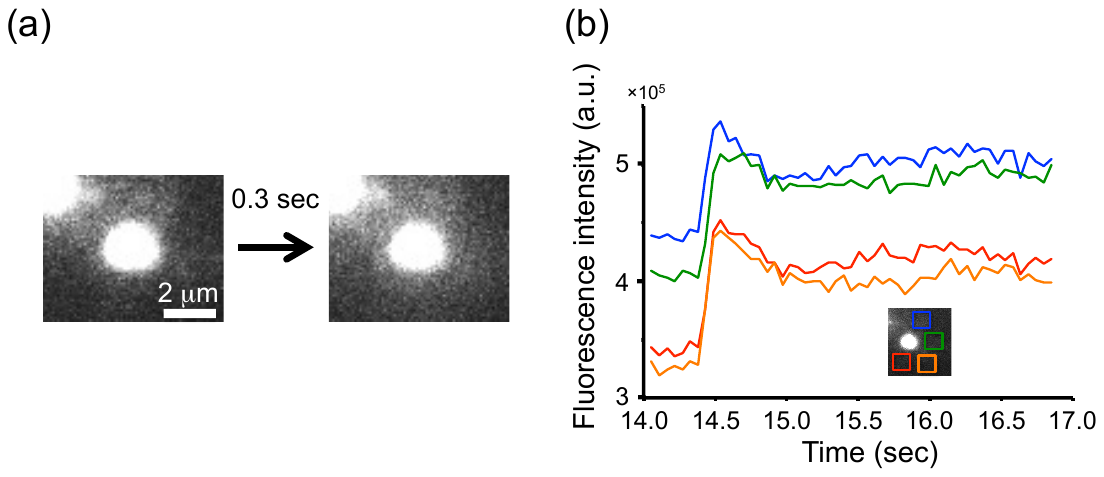

**Supplemental Figure 4.** Leakage of IR700 from vesicles induced by the photoactivation of IR700. (a) The change in fluorescence of IR700 upon photoactivation. (b) Real-time imaging of fluorescence intensity in four color rectangles around a spot of IR700 corresponding to Fig. 4a. The red laser was irradiated from 0 seconds to 17 seconds.

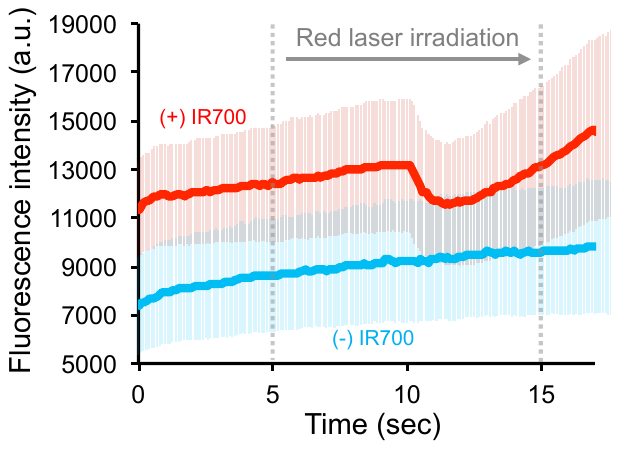

**Supplemental Figure 5.** Change in fluorescence intensity of quantum dots (QDs) in damaged cells. Red and light blue solid lines indicate the fluorescence intensity with or without using IR700 (mean ± SEM). The red laser was irradiated from 5 to 15 seconds.

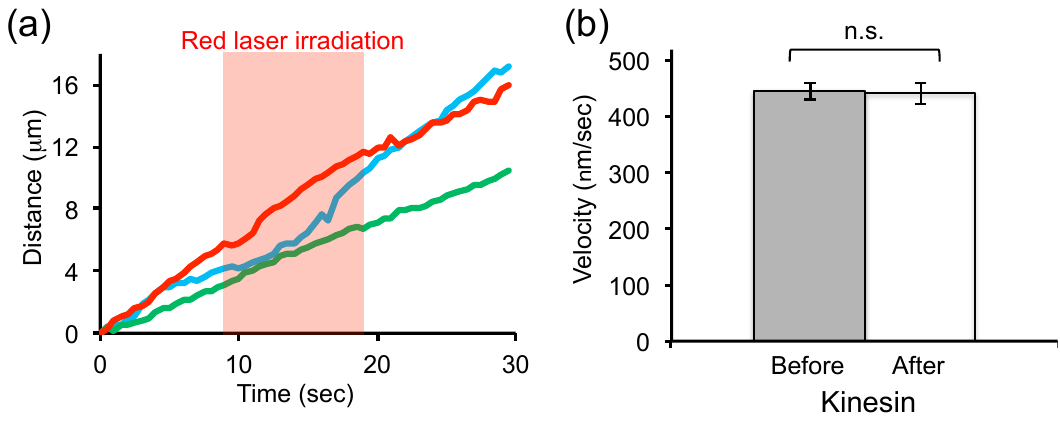

**Supplemental Figure 6.** The effect of the red laser and photoactivation of IR700 with an oxygen scavenging system on the motility of kinesin. (a) Changes in the movement of microtubules without using IR700. Three trajectories indicate the movement of three microtubules. The red laser was irradiated from 8 to 18 seconds. (b) Change in the velocity of microtubules upon photoactivation of IR700 using an oxygen scavenging system (GCOβ). Gray and white bars indicate the velocity before and after red laser irradiation (mean ± SEM), respectively.
